# HVRLocator: A Computationally Efficient Tool for Identifying Hypervariable Regions in 16S rRNA Big Datasets

**DOI:** 10.1101/2025.07.24.666487

**Authors:** Clara Arboleda-Baena, Felipe Borim Correa, Joao Pedro Saraiva, Santiago Castillo, Jonas Coelho Kasmanas, Antonis Chatzinotas, Stephanie D. Jurburg

## Abstract

**Background:** Amplicon sequencing of the 16S rRNA gene is widely used to assess microbial diversity due to its cost-effectiveness and efficiency. However, public 16S rRNA datasets often lack standardized metadata, particularly information on the sequenced hypervariable regions or primers used, which are critical for accurate analysis and data reuse. To address this, we present the HVRLocator, a computational tool that reliably identifies sequenced hypervariable regions, enhancing metadata quality and enabling more robust large-scale microbiome studies.

**Results:** The HVRLocator tool processed samples at an average rate of 0.147 per minute. Validation confirmed 100% accuracy in predicting alignment positions, correctly matching sequences to the expected primer regions based on literature. We demonstrated how to use the tool to select appropriate and comparable sequences for building a global bacterial database from V4 region amplicons of the 16S rRNA gene. Using HVRLocator, we selected 36,217 valid samples out of 45,882 runs, enabling us to identify cases where metadata incorrectly labeled sequences as targeting the V4 region.

**Conclusion:** Even when metadata is available, it can be inaccurate or misleading. HVRLocator offers a reliable and efficient method to identify the exact hypervariable sequenced region, ensuring accurate processing of large-scale 16S rRNA amplicon data. By bypassing inconsistent metadata and literature, it streamlines data curation and enhances the reliability of microbial studies, syntheses, and meta-analyses. Its use is essential for critically evaluating published data and enabling accurate and reproducible research in microbial ecology.

## Background

While the existence of bacteria has been known for over three centuries, the ability to study all individuals in a bacterial community is relatively novel. By extracting and sequencing nucleic acids from hosts or environmental samples, it is now possible to characterize a bacterial community’s taxonomic diversity without the need to cultivate its members. Amplicon sequencing, which focuses the sequencing efforts on a segment of a universal marker gene (typically the 16S rRNA gene for prokaryotes), has emerged as a dominant technique due to its technical ease and low cost. To date, 16S rRNA amplicon sequencing has uncovered the extreme diversity and ubiquity of microbes [1] while revealing avenues for improving human health, agricultural productivity, and sustainability [2]. At the same time, amplicon sequencing data archived in public repositories has grown exponentially [3]. These data are uniform in format, are routinely archived with technical and experimental metadata, and are a rich and growing resource for synthetic and large scale research. However, amplicon sequencing data are archived in their raw format, and the metadata needed for bioinformatics processing is often unavailable or not curated, creating barriers to data reuse [3,4].

Technical metadata are central to sequence data harmonization and reuse as they provide context [5]. Technical choices preceding sequencing, most notably the DNA extraction kit [6], sequencing platform [7], and target amplicon [8] have been shown to affect microbial diversity assessments [9]; however the integration of these data *in light* of processing metadata has received less attention. For example, Abdill and colleagues [10] limited their synthesis to sequences obtained from Illumina technologies to use a unified processing pipeline, but did not enforce consistent amplicon sequence lengths, even though bacterial diversity increases linearly with amplicon length [11].

The 16S rRNA gene contains both highly conserved regions essential for primer design, and hypervariable regions that allow for the phylogenetic identification of microorganisms [12]. Full-length 16S rRNA gene sequences (∼1500 bp) comprise nine hypervariable regions interspersed with nine highly conserved regions [13,14]. Identifying which hypervariable region was targeted for amplification is crucial to fully leverage sequence length and coverage, ultimately determining the efficiency and accuracy of downstream processing and taxonomic classification pipelines. Additionally, in the case of pair-ended reads (e.g., Illumina technologies), the length of the target region informs the minimum read length needed to achieve a successful merger of the pair [15]. The amplified segment may vary in length, directly affecting the diversity detected [11].

At the same time, the choice of the sequenced region can significantly affect the relative abundances of detected organisms. For example, Wasimuddin and colleagues [8] found that, compared to three other primer sets targeting different regions, the primer pair 515F-806R targeting the V4 region of the 16S rRNA gene produced the highest estimates of species richness and diversity across various sample types (including soil, maize roots, cattle rumen, and cattle and human feces) in a model food chain. Similarly, Alberdi and colleagues [16] not only found that one primer set performed better than others, but also demonstrated that the performance of a primer set on real eDNA extracts can differ significantly from predictions based on in silico studies. Consequently, primer choice directly influences the estimation of community composition both within and across compartments and may lead to preferential detection of specific taxa [8]. Crucially, as novel sequencing technologies and platforms have emerged, the length of the target amplicons has varied extensively, from the whole gene. (e.g., nanopore sequencing) to 150 base pair segments (e.g., Illumina HiSeq single-end sequencing), resulting in massive heterogeneity in the length and location of the sequenced regions.

As the sequenced region is a source of technical variation, including this information in the metadata is essential to data reuse, but this information is often missing, vague, or incorrect [17]. This lack of standardized metadata significantly slows down the compilation of large datasets, making it difficult to reprocess sequences collectively and address large-scale questions. Here, we present a computational tool designed to efficiently identify hypervariable regions flanked by primer pairs used in 16S rRNA gene PCR amplification assays. By optimizing computational resources, our approach enables rapid and accurate screening of large datasets, facilitating more comprehensive and scalable microbial diversity analyses.

## Materials and methods

### Design

HVRLocator is a computational tool that identifies which segment of the 16S rRNA gene was sequenced for a given set of amplicon sequencing samples. The full pipeline uses Python programming language and users can access the pipeline either via a singularity container or by installing it locally on their computer. For further details on installation and usage please see: https://github.com/fbcorrea/hvrlocator/.

As input, HVRLocator accepts text file (.txt) lists of accession numbers compliant with International Nucleotide Sequence Collaboration (INSDC) databases including the European Nucleotide Archive (ENA) at the European Bioinformatics Institute, the Sequence Read Archive (SRA) at the National Center for Biotechnology Information, and the DNA Databank of Japan (DDBJ) Sequence Read Archives at the National Institute of Genetics [18] bypassing the need to download the data a priori. HVRLocator also accepts resolved amplicons (e.g., Amplicon Sequence Variants (ASV)) or raw sequencing data in .FASTA or .FASTQ file formats.

For INSDC data, the tool employs a multi-step process. First, it retrieves 1000 reads using fastq-dump from the SRA Toolkit [19] and performs quality control and trimming with fastp [20]. Then, for each sample, the HVRLocator identifies the data type. For paired-end reads, it merges the processed reads using VSEARCH [21], while for single-end reads, it directly converts the trimmed FASTQ to FASTA format. The tool then aligns processed sequences to a reference 16S rRNA gene sequence from an *Escherichia coli* reference genome (J01859.1) using MAFFT [22]. HVRLocator subsequently analyzes the aligned sequences to determine hypervariable regions by identifying the median start and end alignment positions relative to the *E. coli* 16S rRNA gene reference sequence, using established coverage thresholds (default = 0.6) for the conserved and hypervariable regions of the 16S rRNA gene [12,23]. To note, the thresholds can be adjusted by adding the “-t” flag. The output is a tab-separated values (TSV) file containing alignment start and end positions, as well as the boundaries (median and average start and end positions, and minimum start and maximum end positions) of the identified hypervariable regions.

Additionally, we used a Random Forest (RF) model designed to predict the presence of a primer in a given SRA sequencing dataset by analyzing the quality score distribution of the initial subset of reads. To this purpose we selected a curated collection of SRA runs known to possess (882 RUNS) or not sequencing primers (8940 RUNS) (**Supplementary Tables S1 and S2**). For each sample, the first 1,000 reads from each sample were extracted using fastq-dump (NCBI SRA Toolkit, 3.2.1) and two quality score segments from positions 1-5 and 6-10 were calculated. Eight statistical features were obtained-count, mean, median, standard deviation, minimum, maximum, an estimate of skewness (approximated by the 25th percentile), and kurtosis (approximated by the 75th percentile) resulting in 16 features per sample. The model was trained using scikit-learn’s RandomForestClassifier (v1.2.1) with 100 estimators and a fixed random seed (random_state=42), using an 80/20 stratified train-test split. The Random Forest model yielded a precision of 99.96% for the dataset without primers and 100% for the dataset with primers. Recall of the model using the “no-primer” and “primer” dataset was 100% and 99.55%, respectively. A full detail on the model generation including the algorithm, versions and packages is available in the **Supplementary Table S3.**

HVRLocator is available as a singularity container located at https://cloud.sylabs.io/library/jsaraiva/repo/hvrlocator, and can be executed on High Performance Computing (HPC) clusters or cloud computing platforms. Further, Singularity enables the seamless execution of containers without requiring root privileges, maintaining security and reproducibility. The HVRLocator output is a text file (.txt) containing the following columns:

**1. Sample_ID:** Identifier of the processed sample (Run Accession number).
**2. Primer Presence:** Presence or absence of a primer (TRUE/FALSE)
**3. Score Primer Presence:** The associated probability value ranging from 0 to 1.
**4. Min/Max Alignment Start/End:** Minimum (0) and maximum (1540) possible alignment positions along the 16S rRNA gene.
**5. Average Alignment Start/End:** The mean position where reads align to the 16S rRNA gene. The *start* indicates the average starting position, and the *end* indicates the average ending position across all reads in the sample.
**6. Median Alignment Start/End:** The median position where reads align to the 16S rRNA gene. The start indicates the median starting position, and the end indicates the median ending position across all reads in the sample.
**7. Predicted HV region Start/End:** Predicted hypervariable region based on the median alignment start and end positions across all reads, inferred from literature on conserved and hypervariable regions of the 16S rRNA gene (Brosius et al., 1978; Yang et al., 2016).
**8. Coverage based HV region Start/End:** Predicted hypervariable region based on coverage at the start and end positions across all reads.
**9. Coverage HV region Start/End:** Coverage values (0-1) for the “*Coverage based HV region*” start or end position across all reads.
**10. Warnings:** Alerts about low coverage regions. See possible errors and troubleshooting.
**11-19. Cov_V1 to Cov_V9:** Coverage values (0-1) for each HV region.

### Validation

HVRLocator‘s processing stability was calculated by measuring the run time for 1, 10, 100, 1000 and 10000 samples using the same cluster resources: 8 GB of RAM and 4 CPU cores to emulate the standard capabilities of a personal computer.

HVRLocator was validated by analyzing datasets which contained samples sequenced using a) same primer and same sequencing platform, b) the same primer and different sequencing platforms, and c) different primers and the same platform and d) both different primers and sequencing platforms. The first dataset included 17,537 samples from the Earth Microbiome Project (https://earthmicrobiome.org), in which all samples were sequenced using the 515F–806R primer set targeting the V4 region of the 16S rRNA gene and illumina MiSeq sequences [24]. The second dataset is derived from the MiCoDa database Version 1 (https://micoda.idiv.de/) [25], where the samples were sequenced in the same 515-806 region of the 16S SSU rRNA but with a different sequencing platform. For this dataset, we selected 18,426 runs. The third dataset consists of two bioprojects from works that used different primers but were sequenced on the same platform [8,26]. A total of 242 runs were analyzed, and the primer information for each sample was obtained from the metadata archive in NCBI. Finally, for the fourth dataset, we selected 5,308 runs collected from Datathon activities in Latin America [27], which used various sets of primers targeting different regions of the 16S SSU rRNA gene and were sequenced on different platforms. The Bioproject accession numbers used for all datasets are listed in **Supplementary Table S4**.

The results have been computed at the High-Performance Computing (HPC) Cluster EVE, a joint effort of both the Helmholtz Centre for Environmental Research - UFZ (http://www.ufz.de/) and the German Centre for Integrative Biodiversity Research (iDiv) Halle-Jena-Leipzig (http://www.idiv-biodiversity.de/) [28].

## Results

As the number of samples increased, the running time per sample and computational resource usage remained stable (**Figure 1 and Supplementary Table S5**), highlighting the tool’s processing stability and scalability for the analysis of large datasets. The average rate of successfully processed samples per minute is 0.147, calculated across all thresholds and including cases where samples were not processed due to various warnings. Failures were primarily due to runs with fewer than 500 reads (72%), alignment errors (14%), missing FASTQ files (13%), and NCBI portal-related issues (1%). (See *Possible Errors and Troubleshooting* at https://github.com/fbcorrea/hvrlocator).

**Figure 1.**
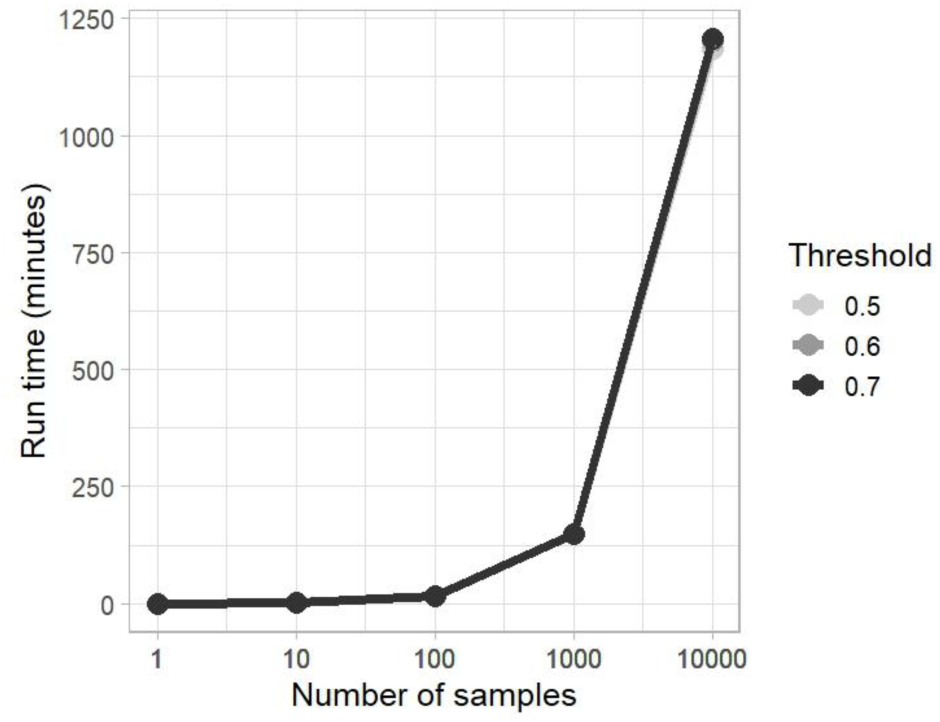
Relationship between the number of samples and run time (in minutes) using 8 GB of RAM and 4 CPU cores. We randomly selected run accession numbers from the Earth Microbiome Project (Dataset 1), MiCoDa V1 (Dataset 2), and Datathon activities (Dataset 4). All samples were downloaded from the NCBI.

Analyzing the alignment positions across different 16S rRNA hypervariable regions and sequencing setups (**Figure 2**), we found that the HVRLocator tool accurately predicted the alignment positions compared to what is expected from the literature associated with each database. For example, sequences from the Earth Microbiome Projects sequences aligned with the V4 hypervariable region of the 16S rRNA gene and had a median sequence start of 532 bp, consistent with the standardized primer set (515F-806R) used in this project. Sequence lengths were highly homogeneous, likely due to the use of the same primer set and sequencing machinery (**Figure 2a**). Out of 16,059 samples processed, 9 did not yield the expected results based on the literature. This highlights the importance of using HVRLocator to critically evaluate published data before conducting analyses, particularly when working with large-scale datasets.

**Figure 2.**
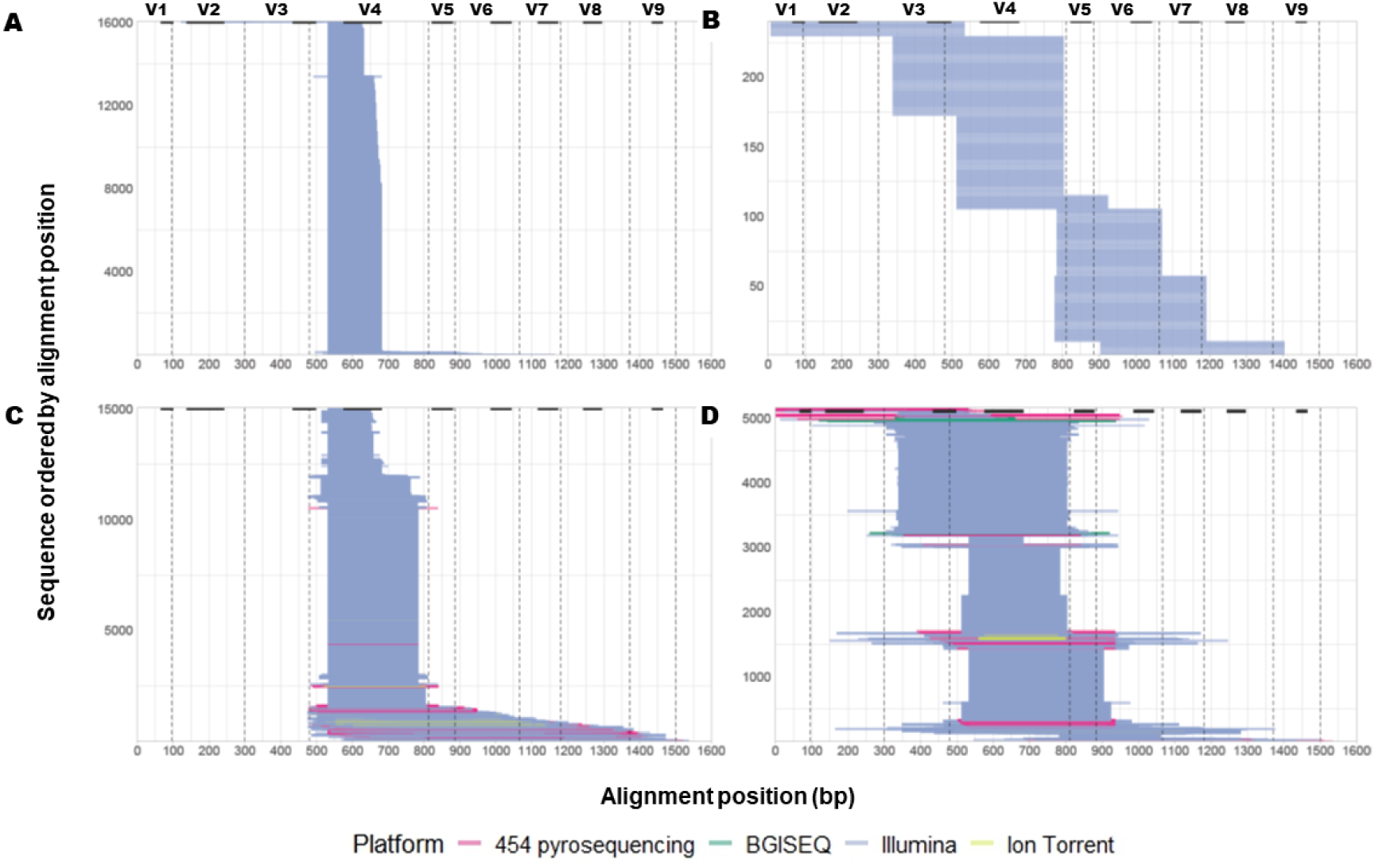
Alignment positions across 16S rRNA regions and sequencing setups. a) Same 16S rRNA region and sequencing setup (N = 16059 runs); b) Different 16S rRNA regions, same sequencing setup (N = 239 runs); c) Same 16S rRNA region, different sequencing setups (N = 15049 runs); d) Different 16S rRNA regions and sequencing setups (N = 5113 runs).The upper part of the figure, along with the dashed lines, indicates the start and end positions of the sequencing setups used to assign specific regions of the 16S rRNA gene (modified from Yang *et al.,* 2016). The hypervariable regions corresponding to each setup are highlighted with bold black bars (Brosius *et al.,* 1978).

The datasets from Wasimuddin et al. and Varliero et al. [8,26], which employed different primers but were sequenced on the same platform, allowed us to confirm that HVRLocator correctly matched the sequences to the corresponding primer regions used during sample sequencing with 100% accuracy (**Figure 2b**). For the dataset that targeted the same region with different sequencing platforms (**Figure 2c**), HVRLocator also accurately showed that 90% of the sequences start in the V4 region of the 16S rRNA gene, but indicated a more heterogeneous alignment position start and sequence lengths, consistent with our expectation. HVRLocator also correctly identified the different regions sequenced by the dataset targeting different 16S rRNA regions with the same sequencing platform (**Figure 2c).** Of the 16,771 samples processed, 1,712 (10%) did not produce results consistent with the literature. This underscores the value of validating literature data with HVRLocator before analysis. Finally, for the panel visualizing the results of runs that used both different 16S rRNA regions and sequencing setups (**Figure 2d**), HVRLocator accurately assigned the alignment positions for this diverse dataset.

An important outcome of the HVRLocator tool is its ability to identify problematic sequences. For example, in **Figures 2c and 2d**, we observed sequences with abnormally long lengths, exceeding 600 bp, which is beyond the typical output of the Illumina sequencing platform. Upon reviewing these sequences, BioProject by BioProject, we found that either the sequencing platform was incorrectly annotated in the metadata (NCBI or the associated publication), or the sequences did not correspond to the 16S rRNA gene but instead to ITS or the nifH gene. This underscores the importance of using HVRLocator as a curation tool for large datasets, where human errors in annotation can significantly impact downstream analysis. Additionally, it is crucial to apply this prior to merging sequences and performing taxonomic analysis, even in datasets that have been previously used, in order to detect potential issues.

Alternatively, the predicted hypervariable region based on coverage for the second dataset (MiCoDa Database Version 1) is shown in **Figure 3**. These samples were sequenced for the V4 region of the 16S SSU rRNA gene but using a different sequencing platform. The results confirm that most sequences covered the V4 region (89%), as indicated by both the Median Alignment Start and the Coverage-Based HV Region Start (**Figure 2c and Figure 3**, respectively). As expected, the Median Alignment End varies across projects and sequencing platforms, showing differences in coverage between regions V4 and V9 as well.

**Figure 3.**
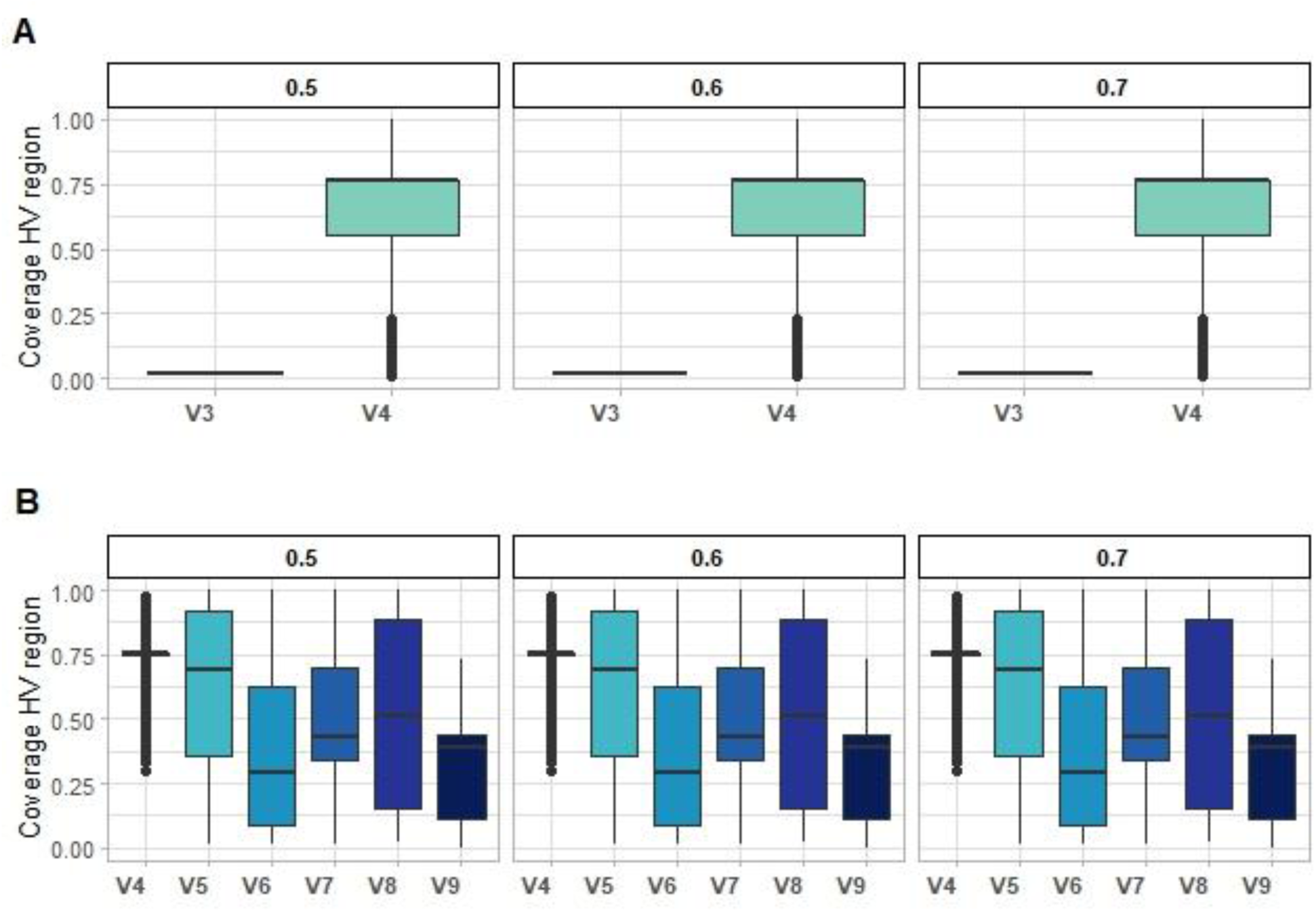
Differences in 16S rRNA gene coverage using the same primer set (Primer 515R-806R for V4 region) but different sequencing setups. A) Predicted 16S rRNA region coverage start, B) Predicted 16s rRNA region coverage end.

### Case study: 45,882 amplicon sequencing samples for the development of a global V4 16S rRNA gene database

We presented an example of how to use the tool to select correct and comparable sequences for constructing a global bacterial database based on amplicon sequences from the V4 region of the 16S rRNA gene. We used data identified as potentially applicable to the MiCoDa v2 (https://micoda.idiv.de/) [25], which included samples sequenced from start position 515 bp of the 16S rRNA gene, the same region using the Earth Microbiome Project primers (525-806R) [1]. Our input dataset included 45,882 runs spanning a wide variety of matrices (e.g., soil, host-associated, and water), sequenced with different primer sets and sequencing platforms and collected through an extensive literature search prioritizing meta-analyses, large amplicon research consortia, and Datathon activities [27] (**Supplementary Table S6**). Using SRA’s sequence-associated metadata, we selected only amplicon sequences (i.e., excluding WGA, WGS, Tn-Seq, miRNA-Seq, POOLCLONE, RNA-Seq, etc).

HVRLocator was run on 45,882 amplicon sequencing samples, processing approximately one sample every 0.15 minutes, using 8 GB of RAM and 4 CPU cores. A total of 42,166 samples were processed successfully, while 3,716 samples failed to process and generated warnings, mainly due to runs with fewer than 500 reads (66%), missing FASTQ files (21%), alignment errors (13%), and other issues related to the NCBI portal (1%). (See *Possible Errors and Troubleshooting* at https://github.com/fbcorrea/hvrlocator).

HVRLocator identified a diverse range of sequences with varying hypervariable region start and end positions, coverage, and lengths **(Figure 4a)**. The output showed that 1,532 samples had a true presence of primers, while 40,634 did not.

**Figure 4.**
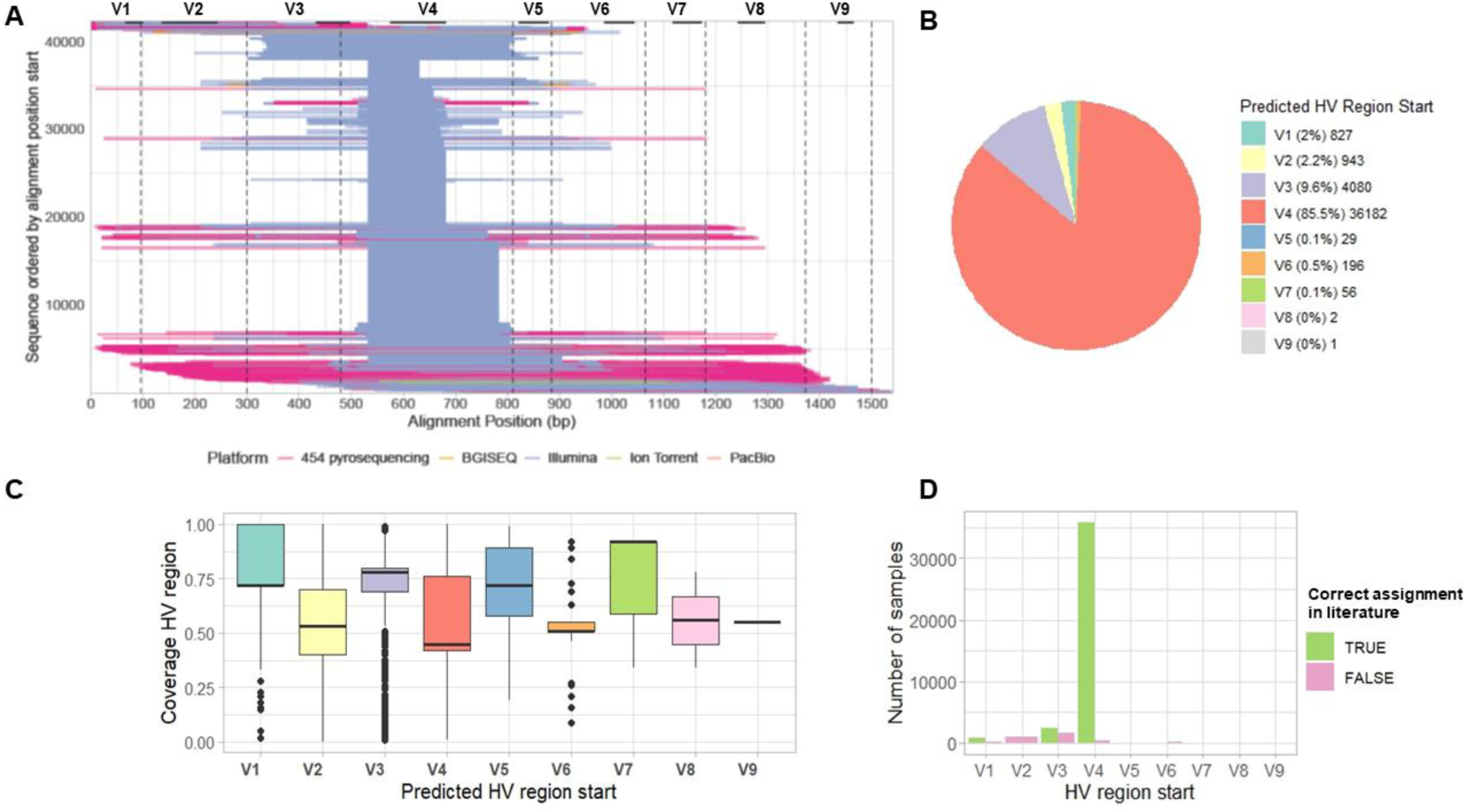
Application of HVRLocator for the selection of V4 16S rRNA gene amplicon sequences from MiCoDa V2. A) Alignment start positions across the 16S rRNA gene for the 42,316 runs analyzed, B) Percentage of samples retained for downstream analyses after applying the HVRLocator tool, C) Variation in gene coverage across sequences, D) Number of samples in which primer assignment matched the metadata.

The average alignment column showed that 77.5% of the samples (32,677) were within the expected range for the V4 region start position 501-550 pb (**Supplementary Table S7a**), while the end position of these specific samples varied from 601 to 1500 pb; 45.8% of the sequences ended from 651-700 pb, whereas 30.8% from 751-800 pb (**Supplementary Table S7b**). Although this criterion provided a basis for selecting sequences within the desired range for the target database, we advocated for an alternative selection based on the Median Alignment Start/End positions, as detailed in the following discussion.

The results of the median alignment columns showed for the start position 500-550 pb a total of 34,474 samples (81.8%) (**Supplementary Table S7c**). These have an end position from 601 to 1550 pb where the higher percentage are for sequences that ended from 651-700 pb (47.8%) and from 751-800 pb (31.3%) (**Supplementary Table S7d**). With this tool, we provide two metrics: the average and the median alignment positions, for both the start and end of sequences. However, our validation showed that the median is more reliable, as low-quality sequences within a sample can skew the mean and lead to incorrect identification of the hypervariable region. For this reason, we report both metrics but recommend prioritizing the median when deciding whether to retain or discard sequences during downstream processing.

Based on the predicted HV region start, 85.9% (36,217) of the 42,166 samples began at the V4 region of the 16S rRNA gene (**Figure 4b**). The next most common starting regions were V3 (9.7%), V2 (2.3%), and V1 (1.5%). For our analyses, we retained the 36,217 samples that had a median starting point in the V4 region. In contrast, the predicted hypervariable region end for the majority of samples also corresponded to the V4 region (85.3%), followed by V6 (6.5%), V8 (3.4%), V5 (1.6%), and V7 (1.4%) (**Supplementary Table S7e**). However, one output indicated the start and end positions of the sequences, while the other provided the predicted hypervariable region based on coverage at those positions across all reads. In the output column for the coverage-based HV region start (**Figure 4c**), 36,130 samples began in the V4 region, with a median coverage of 0.57 (**Supplementary Table S7f**). Among these samples, the coverage-based HV region end corresponded to the V4 region in 33,022 samples (91.4%), V5 in 2079 samples (5.7%), V6 in 505 samples (1.4%), V7 in 287 samples (0.8%), V8 in 236 samples (0.6%), and V9 in 1 sample (**Supplementary Table S7g**). It is important to note that, in general, the warnings indicate that the coverage of a predicted hypervariable region is below the chosen threshold, which is 0.6 in this case. Each warning corresponds to a predicted hypervariable region falling below this threshold (**Supplementary Table S7h**).

Finally, the HVRLocator enabled us to select 36,217 samples from a total of 45,882 runs. Among the selected samples, 382 had incorrect primer annotations, either in the NCBI metadata or in the associated publications. For all sequences, we cross-checked the reported primer information either from the metadata of the research articles or NCBI records against the region predicted by HVRLocator (**Figure 4d**). This allowed us to quantify the number of samples where, despite being labeled as targeting the V4 hypervariable region in the metadata, the actual sequenced region was incorrectly assigned. These findings highlight that even when metadata is available, it may be inaccurate or misleading. This underscores the importance of tools like HVRLocator, which accurately and efficiently identify the sequenced region. Such precision is critical for the reliable processing of large datasets, particularly in large-scale studies, synthesis efforts, and metanalyses.

## Discussion

The INSDC now contains over 32 million next-generation sequencing runs and 4.76 billion assembled sequences [29], a growing resource for large-scale analyses to address global questions through synthesis and meta-analytical research. However, efforts of sequence data archiving are undermined by the lack of available metadata [16], especially, as these metadata are crucial to data processing and high quality data are relatively sparse [3], which makes the data reuse process curation-intensive, inefficient, and could introduce errors. In the case of 16S rRNA amplicon sequences, information about the gene region sequenced is essential to the bioinformatics processing of the sequence data, and for the statistical analyses (i.e., as a random effect in a hierarchical model) and data interpretation downstream. To facilitate the reuse of bacterial amplicon sequence data, we developed HVRLocator, a publicly available tool which efficiently identifies the exact region sequenced by a set of 16S rRNA sequences.

Extensive validation of HVRLocator also highlights its potential for application towards data reuse.

Given the ubiquity of bacteria and their relevance to their environments a wide range of disciplines employ 16S rRNA amplicon sequencing, and contribute data to INSDC archives in this process. Given the lack of curation of INSDC metadata, information derived from peer-reviewed literature has been proposed as a central source of technical metadata that can enrich existing datasets [30]. The diversity of disciplines also results in different degrees of resolution in the technical metadata provided for the sequence data, and may lack the resolution necessary for improved bioinformatics process on, or even introduce errors. Here, HVRLocator serves to bypass the need to return to the original literature to obtain necessary processing metadata, to obtain higher resolution information (i.e., the exact start and end sequence positions instead of the general region sequenced), and to correct potential errors that might be present in the literature-derived metadata.

Knowing which region was sequenced is crucial, as species identification depends heavily on the targeted region [11]. Without this information, making decisions about which sequences to include in a large-scale study, as well as comparing datasets, becomes difficult and may lead to biased or incompatible results. Additionally, from an ecological perspective, the ability to consistently target the same genetic region across different studies brings us closer to achieving a macroecological understanding of microbial communities [8,31].

Nevertheless, since this problem depends on how users archive data, it is important to identify strategies to improve and accelerate data reuse in a more systematic and efficient manner, both in terms of accuracy and time, to address large-scale research questions. In this vein, the use of HVRLocator may greatly accelerate the identification of sequences of the correct gene and the correct region of the gene, even when metadata is incomplete, and links to the associated peer reviewed literature are unavailable.

## Availability of supporting source code and requirements

**Project name: HVRLocator**

**Project home page:** https://github.com/fbcorrea/HVRLocator

**Operating system(s):** Linux OS

**Programming language:** Python 3.9

**Other requirements:** Singularity container platform

**License:** GNU GPL-3.0 **RRID:** Not applicable **bio.tools ID:** Not applicable

## Additional files

**Supplementary Table S1:** Run accession list with primers used for training and testing the Random Forest (RF) model.

**Supplementary Table S2:** Run accession list without primers used for training and testing the Random Forest (RF) model.

**Supplementary Table S3:** Random Forest Model for Primer Presence Prediction.

**Supplementary Table S4:** Bioprojects included in the validation process.

**Supplementary Table S5:** Number of samples per dataset and run time (in minutes) using 8 GB of RAM and 4 CPU cores.

**Supplementary Table S6:** Run accession list for case study

**Supplementary Table S7:** Case study output.

**Supplementary Figure S1:** Variation in gene coverage across sequences and number of samples in which primer assignment matched the metadata.

## Data Availability

The public datasets used in this paper can be found in **Supplementary Table S3** and Table S5.

## Abbreviations

ASV: Amplicon Sequence Variants
DDBJ: DNA Databank of Japan
DRA: Sequence Read Archives
EBI: European Bioinformatics Institute (EBI)
ENA: European Nucleotide Agency
HPC: High Performance Computing
INSDC: International Nucleotide Sequence Database Collaboration
NCBI: National Center for Biotechnology Information
NIG: National Institute of Genetics
SRA: Sequence Read Archive
TSV: Tab-separated values
UFZ: Helmholtz Center for Environmental Research.

## Acknowledgments

We would like to thank the administration and support staff of EVE who keep the system running and support us with our scientific computing needs: Toni Harzendorf, Mark Fliak and Conrad Ostertag from UFZ, and Christian Krause from iDiv. Also, we would like to thank Marten Winter at the Synthesis Centre for Biodiversity Sciences (sDiv) at the German Centre for Integrative Biodiversity Research (iDiv).

## Author Contributions

Conceptualization: SJ, FBC, CAB; funding acquisition: SJ; methodology: SJ, FBC, CAB, JPS, SC; software: FBC, JPS; writing—original draft: SJ, FBC, CAB, JPS; writing—review and editing: all authors. All authors read and approved the final manuscript.

## Funding

sIBTEDS project (Illuminating Blindspots Through Equitable Data Reuse practices in the Global South) from the German Centre for Integrative Biodiversity Research (iDiv).

## Competing Interests

The authors declare that they have no competing interests

## SUPPLEMENTARY MATERIAL

**Table S4.**
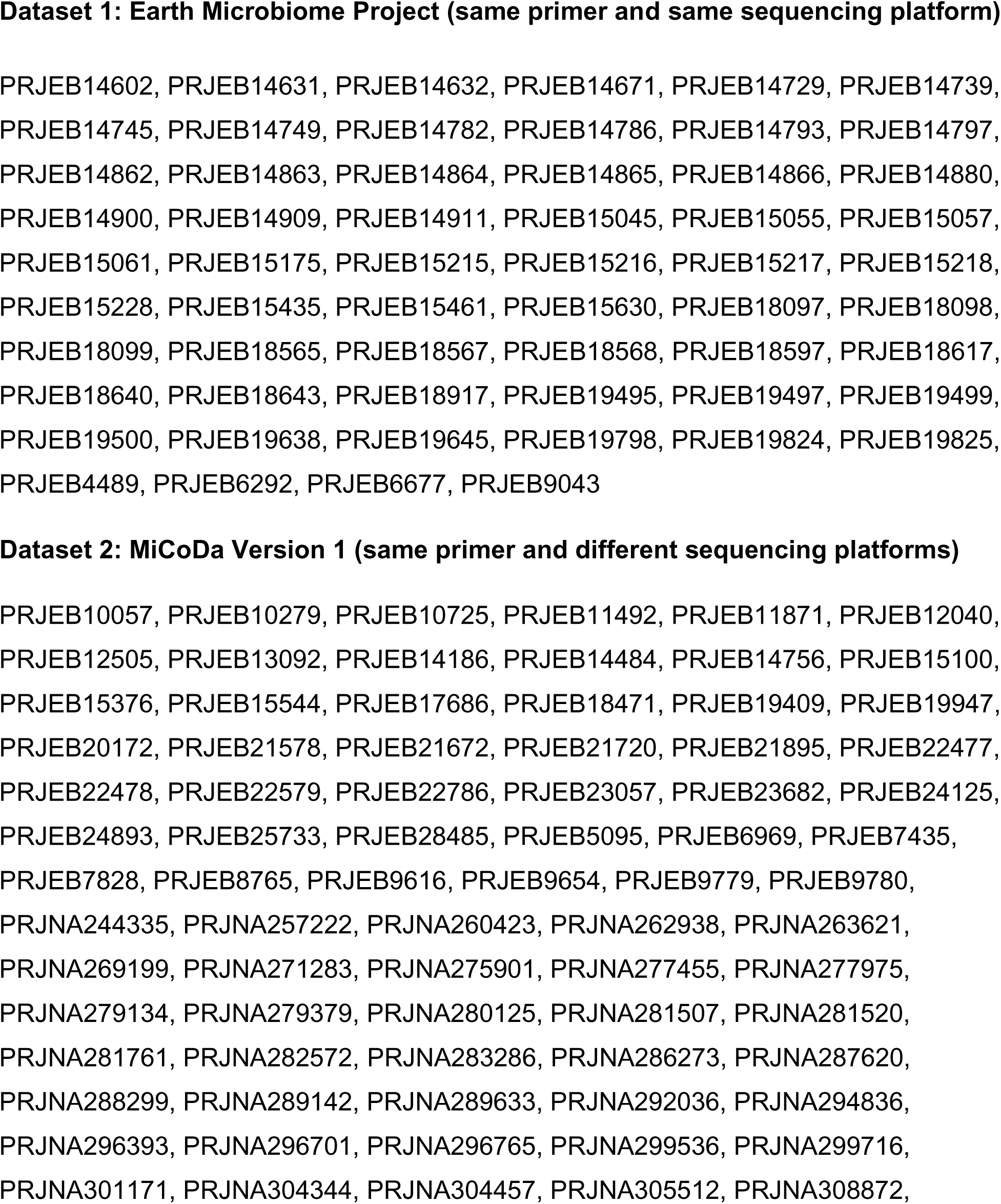
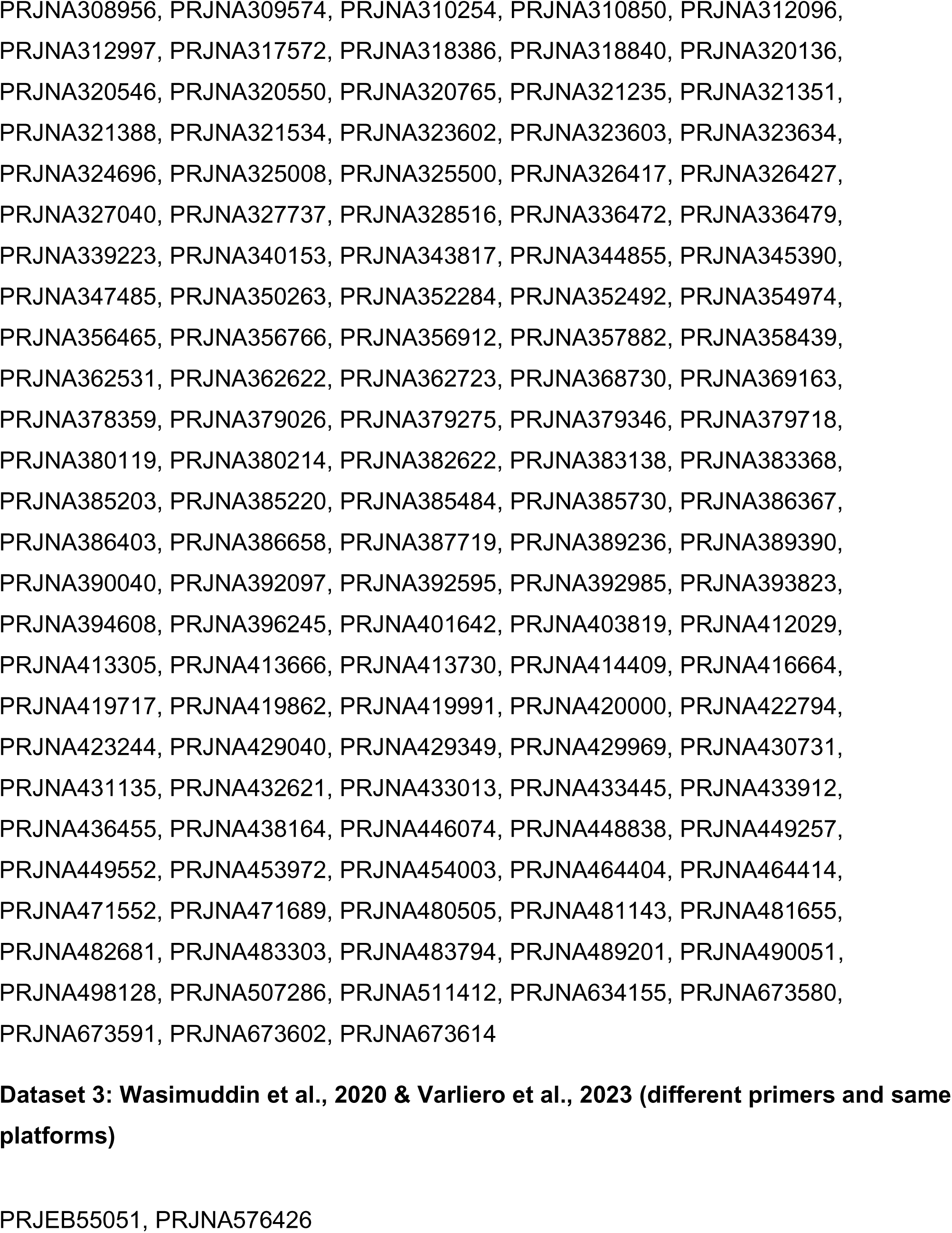
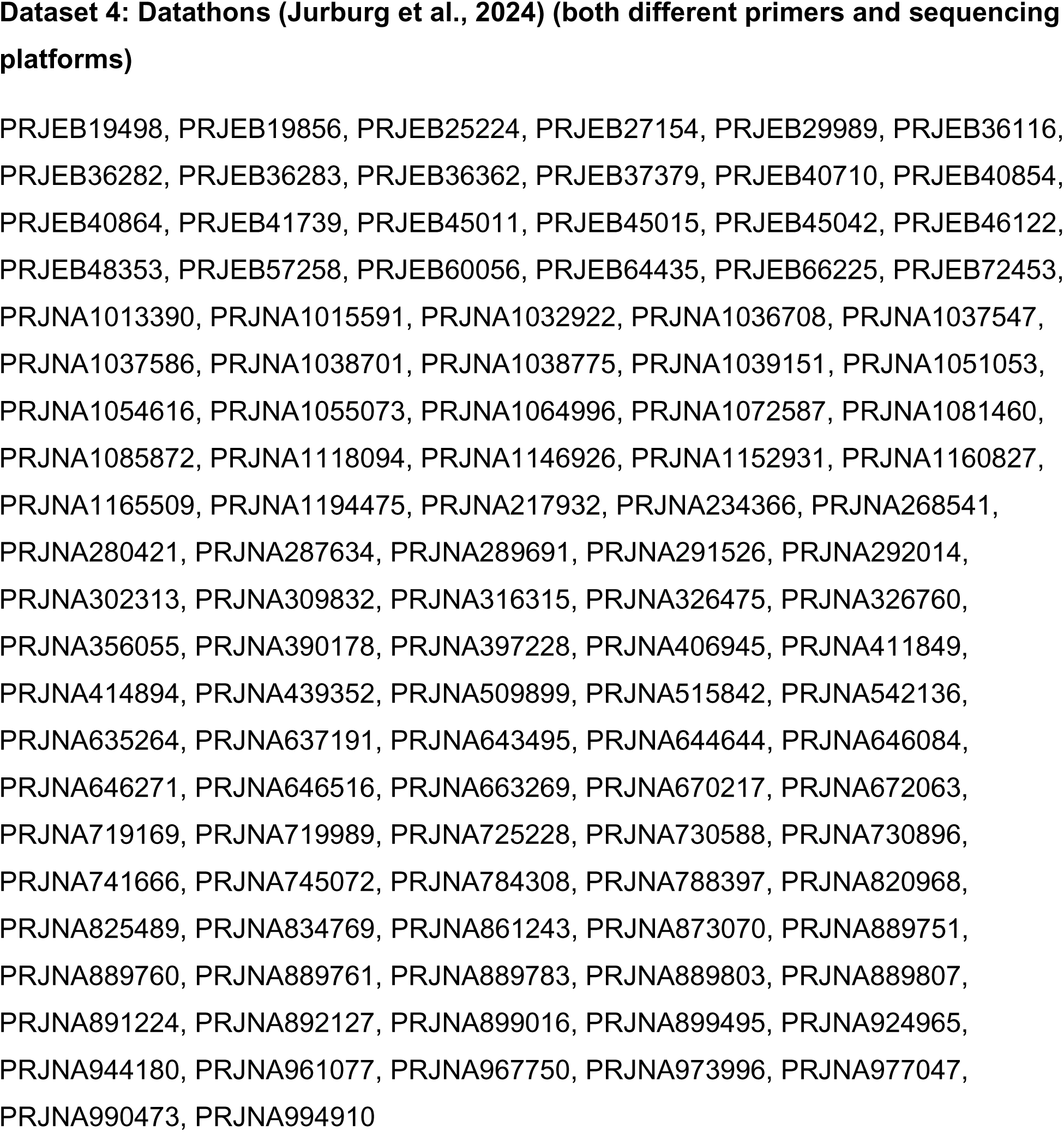
Bioprojects Included in the Validation Process.

**Table S5:** Number of samples per dataset and run time (in minutes) using 8 GB of RAM and 4 CPU cores. We selected run accession numbers from the Earth Microbiome Project (Dataset 1), MiCoDa V1 (Dataset 2), and Datathon activities (Dataset 4). All samples were downloaded from the NCBI.

| **Dataset** | **Threshold** | **Duration**<br>**(minute)** | **Total Samples**<br>**(Run Accession**<br>**Numbers)** | **Samples processed**<br>**successfully** | **Samples Not**<br>**Processed**<br>**(Warnings)** | **Samples processed**<br>**per minute** |
| --- | --- | --- | --- | --- | --- | --- |
| **1** | **0.5** | 2078.26 | 17537 | 16059 | 1478 | 0.1185 |
| **2** | **0.5** | 2746.85 | 18426 | 16771 | 1655 | 0.1491 |
| **4** | **0.5** | 922.38 | 5308 | 5163 | 145 | 0.1738 |
| **1** | **0.6** | 2106.22 | 17537 | 16059 | 1478 | 0.1201 |
| **2** | **0.6** | 2746.85 | 18426 | 16771 | 1655 | 0.1491 |
| **4** | **0.6** | 904.98 | 5308 | 5163 | 145 | 0.1705 |
| **1** | **0.7** | 2114.30 | 17537 | 16059 | 1478 | 0.1206 |
| **2** | **0.7** | 2759.60 | 18426 | 16771 | 1655 | 0.1498 |
| **4** | **0.7** | 917.05 | 5308 | 5163 | 145 | 0.1728 |

**Table S7:** Case study outputs.

**A. Average Alignment Start/End**
| Average alignment start |  |  | Average alignment end |  |  |
| --- | --- | --- | --- | --- | --- |
| Range | Number of Samples | Percentage | Range | Number of Samples | Percentage |
| 501-550 | 32677 | 77.50 | 651-700 | 15414 | 36.56 |
| 301-350 | 2528 | 6.00 | 751-800 | 10908 | 25.87 |
| 551-600 | 1707 | 4.05 | 801-850 | 5595 | 13.27 |
| 451-500 | 1033 | 2.45 | 601-650 | 2773 | 6.58 |
| 351-400 | 772 | 1.83 | 901-950 | 1841 | 4.37 |
| 601-650 | 680 | 1.61 | 701-750 | 854 | 2.03 |
| 401-450 | 398 | 0.94 | 1051-1100 | 607 | 1.44 |
| 1-50 | 387 | 0.92 | 851-900 | 525 | 1.25 |
| 651-700 | 367 | 0.87 | 1301-1350 | 523 | 1.24 |
| 201-250 | 353 | 0.84 | 951-1000 | 513 | 1.22 |
| 151-200 | 279 | 0.66 | 1001-1050 | 506 | 1.20 |
| 251-300 | 229 | 0.54 | 1251-1300 | 439 | 1.04 |
| 751-800 | 205 | 0.49 | 1101-1150 | 364 | 0.86 |
| 701-750 | 118 | 0.28 | 1151-1200 | 346 | 0.82 |
| 951-1000 | 113 | 0.27 | 301-350 | 340 | 0.81 |
| 801-850 | 94 | 0.22 | 1201-1250 | 303 | 0.72 |
| 101-150 | 86 | 0.20 | 501-550 | 121 | 0.29 |
| -49-0 | 43 | 0.10 | 1351-1400 | 61 | 0.14 |
| 51-100 | 40 | 0.09 | 551-600 | 57 | 0.14 |
| 851-900 | 26 | 0.06 | 1401-1450 | 49 | 0.12 |
| 901-950 | 16 | 0.04 | 451-500 | 17 | 0.04 |
| 1101-1150 | 9 | 0.02 | 1451-1500 | 6 | 0.01 |
| 1001-1050 | 3 | 0.01 | 201-250 | 1 | 0.00 |
| 1051-1100 | 2 | 0.00 | 251-300 | 1 | 0.00 |
| 1151-1200 | 1 | 0.00 | 351-400 | 1 | 0.00 |
|  |  |  | 401-450 | 1 | 0.00 |
| **TOTAL** | **42166** | **100.00** | **TOTAL** | **42166** | **100.00** |

**B. Average Alignment End of the previous 32677 sequences with average alignment start position at 501-550 pb.**
| Average alignment end |  |  |
| --- | --- | --- |
| Range | Number of Samples | Percentage |
| 651-700 | 14975 | 45.83 |
| 751-800 | 10081 | 30.85 |
| 601-650 | 2712 | 8.30 |
| 801-850 | 2651 | 8.11 |
| 901-950 | 1287 | 3.94 |
| 701-750 | 372 | 1.14 |
| 1001-1050 | 155 | 0.47 |
| 851-900 | 126 | 0.39 |
| 951-1000 | 82 | 0.25 |
| 1051-1100 | 68 | 0.21 |
| 1201-1250 | 46 | 0.14 |
| 1151-1200 | 42 | 0.13 |
| 1251-1300 | 27 | 0.08 |
| 1301-1350 | 20 | 0.06 |
| 1101-1150 | 19 | 0.06 |
| 1351-1400 | 10 | 0.03 |
| 1401-1450 | 2 | 0.01 |
| 1451-1500 | 2 | 0.01 |
| **TOTAL** | **32677** | **100.00** |

C. Median Alignment Start/End
| Median alignment_start |  |  | Median alignment end |  |  |
| --- | --- | --- | --- | --- | --- |
| Range | Number of Samples | Percentage | Range | Number of Samples | Percentage |
| 501-550 | 34474 | 81.76 | 651-700 | 16736 | 39.69 |
| 301-350 | 2699 | 6.40 | 751-800 | 11041 | 26.18 |
| 451-500 | 616 | 1.46 | 801-850 | 5300 | 12.57 |
| 351-400 | 532 | 1.26 | 601-650 | 2921 | 6.93 |
| 1-50 | 476 | 1.13 | 901-950 | 1863 | 4.42 |
| 401-450 | 455 | 1.08 | 1351-1400 | 783 | 1.86 |
| 601-650 | 439 | 1.04 | 851-900 | 493 | 1.17 |
| 551-600 | 374 | 0.89 | 1101-1150 | 350 | 0.83 |
| 151-200 | 323 | 0.77 | 951-1000 | 349 | 0.83 |
| 201-250 | 301 | 0.71 | 301-350 | 345 | 0.82 |
| 701-750 | 252 | 0.60 | 1201-1250 | 304 | 0.72 |
| 101-150 | 215 | 0.51 | 1051-1100 | 272 | 0.65 |
| 751-800 | 213 | 0.51 | 1151-1200 | 269 | 0.64 |
| 651-700 | 176 | 0.42 | 1301-1350 | 234 | 0.55 |
| 951-1000 | 154 | 0.37 | 1001-1050 | 212 | 0.50 |
| 251-300 | 103 | 0.24 | 1251-1300 | 170 | 0.40 |
| 801-850 | 98 | 0.23 | 501-550 | 155 | 0.37 |
| 51-100 | 92 | 0.22 | 701-750 | 111 | 0.26 |
| -49-0 | 67 | 0.16 | 1401-1450 | 93 | 0.22 |
| 1051-1100 | 37 | 0.09 | 1451-1500 | 79 | 0.19 |
| 1001-1050 | 19 | 0.05 | 551-600 | 31 | 0.07 |
| 1101-1150 | 19 | 0.05 | 351-400 | 18 | 0.04 |
| 901-950 | 18 | 0.04 | 1501-1550 | 15 | 0.04 |
| 851-900 | 11 | 0.03 | 401-450 | 13 | 0.03 |
| 1151-1200 | 1 | 0.00 | 451-500 | 6 | 0.01 |
| 1301-1350 | 1 | 0.00 | 201-250 | 2 | 0.00 |
| 1401-1450 | 1 | 0.00 | 151-200 | 1 | 0.00 |
| **TOTAL** | **42166** | **100** | **TOTAL** | **42166** | **100** |

D. Median Alignment End of the previous 34474 sequences with average alignment start position at 501-550 pb.
| Median alignment end |  |  |
| --- | --- | --- |
| Range | Number of Samples | Percentage |
| 651-700 | 16483 | 47.81 |
| 751-800 | 10782 | 31.28 |
| 601-650 | 2828 | 8.20 |
| 801-850 | 2478 | 7.19 |
| 901-950 | 1418 | 4.11 |
| 851-900 | 178 | 0.52 |
| 701-750 | 72 | 0.21 |
| 1001-1050 | 43 | 0.12 |
| 1151-1200 | 40 | 0.12 |
| 951-1000 | 37 | 0.11 |
| 1201-1250 | 28 | 0.08 |
| 1351-1400 | 26 | 0.08 |
| 1051-1100 | 20 | 0.06 |
| 1101-1150 | 17 | 0.05 |
| 1301-1350 | 11 | 0.03 |
| 1251-1300 | 6 | 0.02 |
| 1401-1450 | 3 | 0.01 |
| 1451-1500 | 3 | 0.01 |
| 1501-1550 | 1 | 0.00 |
| **TOTAL** | **34474** | **100** |

E. Predicted HV region start/end
| Predicted HV region start |  |  | Predicted HV region end |  |  |
| --- | --- | --- | --- | --- | --- |
| HV region | Number of samples | % | HV region | Number of samples | % |
| V1 | 626 | 1.48 | V1 | 0 | 0.00 |
| V2 | 951 | 2.26 | V2 | 3 | 0.01 |
| V3 | 4088 | 9.70 | V3 | 380 | 0.90 |
| V4 | 36217 | 85.89 | V4 | 35959 | 85.28 |
| V5 | 29 | 0.07 | V5 | 692 | 1.64 |
| V6 | 196 | 0.46 | V6 | 2733 | 6.48 |
| V7 | 56 | 0.13 | V7 | 578 | 1.37 |
| V8 | 2 | 0.00 | V8 | 1446 | 3.43 |
| V9 | 1 | 0.00 | V9 | 375 | 0.89 |
| **TOTAL** | **42166** | **100.00** | **TOTAL** | **42166** | **100.00** |

F. Coverage based HV region Start/End
| Coverage based HV region start |  |  | Coverage based HV region end |  |  |
| --- | --- | --- | --- | --- | --- |
| HV region | Number of samples | % | HV region | Number of samples | % |
| V1 | 517 | 1.23 | V1 | 2 | 0.00 |
| V2 | 496 | 1.18 | V2 | 363 | 0.86 |
| V3 | 3759 | 8.91 | V3 | 478 | 1.13 |
| V4 | 36130 | 85.69 | V4 | 35836 | 84.99 |
| V5 | 1004 | 2.38 | V5 | 2600 | 6.17 |
| V6 | 61 | 0.14 | V6 | 865 | 2.05 |
| V7 | 178 | 0.42 | V7 | 803 | 1.90 |
| V8 | 19 | 0.05 | V8 | 1206 | 2.86 |
| V9 | 2 | 0.00 | V9 | 13 | 0.03 |
| **TOTAL** | **42166** | **100.00** | **TOTAL** | **42166** | **100.00** |

G. Coverage based HV region End of the previous 36130 sequences with average alignment start position at 501-550 pb.
| Coverage based HV region end |  |  |
| --- | --- | --- |
| HV region | Number of samples | % |
| V4 | 33022 | 91.40 |
| V5 | 2079 | 5.75 |
| V6 | 505 | 1.40 |
| V7 | 287 | 0.79 |
| V8 | 236 | 0.65 |
| V9 | 1 | 0.00 |
| **TOTAL** | **36130** | **100.00** |

H. Mean coverage associated with the most frequent warning during the samples processing.
| Average of coverage HV region start |  |  |  |  |  |  |  |  |  |
| --- | --- | --- | --- | --- | --- | --- | --- | --- | --- |
| Warnings | Cov_V1 | Cov_V2 | Cov_V3 | Cov_V4 | Cov_V5 | Cov_V6 | Cov_V7 | Cov_V8 | Cov_V9 |
| V1 and V2 below threshold of .6 | 0.45 | 0.42 |  |  |  |  |  |  |  |
| V1 and V3 below threshold of .6 | 0.36 | 1.00 | 0.35 |  |  |  |  |  |  |
| V1 and V4 below threshold of .6 | 0.34 | 1.00 | 1.00 | 0.35 |  |  |  |  |  |
| V1 and V5 below threshold of .6 | 0.37 | 1.00 | 1.00 | 1.00 | 0.31 |  |  |  |  |
| V1 and V6 below threshold of .6 | 0.30 | 1.00 | 1.00 | 1.00 | 1.00 | 0.33 |  |  |  |
| V1 and V8 below threshold of .6 | 0.41 | 1.00 | 1.00 | 1.00 | 1.00 | 1.00 | 1.00 | 0.23 |  |
| V1 below threshold of .6 | 0.39 | 1.00 | 0.95 | 0.73 | 0.65 | 0.65 | 0.65 | 0.58 |  |
| V2 and V3 below threshold of .6 |  | 0.43 | 0.26 |  |  |  |  |  |  |
| V2 and V4 below threshold of .6 |  | 0.38 | 1.00 | 0.45 |  |  |  |  |  |
| V2 and V5 below threshold of .6 |  | 0.21 | 1.00 | 1.00 | 0.28 |  |  |  |  |
| V2 and V6 below threshold of .6 |  | 0.25 | 1.00 | 1.00 | 1.00 | 0.26 |  |  |  |
| V2 and V7 below threshold of .6 |  | 0.28 | 1.00 | 1.00 | 1.00 | 1.00 | 0.32 |  |  |
| V2 and V8 below threshold of .6 |  | 0.39 | 1.00 | 1.00 | 1.00 | 1.00 | 1.00 | 0.27 |  |
| V2 and V9 below threshold of .6 |  | 0.44 | 1.00 | 1.00 | 1.00 | 1.00 | 1.00 | 1.00 | 0.47 |
| V2 below threshold of .6 | 0.28 | 0.47 | 1.00 | 0.99 | 0.91 | 0.88 | 0.78 | 0.67 |  |
| V3 and V4 below threshold of .6 |  |  | 0.40 | 0.38 |  |  |  |  |  |
| V3 and V5 below threshold of .6 |  |  | 0.28 | 1.00 | 0.43 |  |  |  |  |
| V3 and V6 below threshold of .6 |  |  | 0.17 | 1.00 | 1.00 | 0.39 |  |  |  |
| V3 and V7 below threshold of .6 |  |  | 0.37 | 1.00 | 1.00 | 1.00 | 0.37 |  |  |
| V3 and V8 below threshold of .6 |  |  | 0.35 | 1.00 | 1.00 | 1.00 | 1.00 | 0.29 |  |
| V3 and V9 below threshold of .6 |  |  | 0.19 | 1.00 | 1.00 | 1.00 | 1.00 | 1.00 | 0.16 |
| V3 below threshold of .6 | 0.35 | 0.48 | 0.21 | 0.56 | 0.39 | 0.38 | 0.11 | 0.73 |  |
| V4 and V5 below threshold of .6 |  |  |  | 0.46 | 0.43 |  |  |  |  |
| V4 and V6 below threshold of .6 |  |  |  | 0.39 | 1.00 | 0.14 |  |  |  |
| V4 and V7 below threshold of .6 |  |  |  | 0.28 | 1.00 | 1.00 | 0.39 |  |  |
| V4 and V8 below threshold of .6 |  |  |  | 0.18 | 1.00 | 1.00 | 1.00 | 0.18 |  |
| V4 and V9 below threshold of .6 |  |  |  | 0.26 | 1.00 | 1.00 | 1.00 | 1.00 | 0.23 |
| V4 below threshold of .6 | 0.35 | 0.47 | 0.16 | 0.41 | 0.20 | 0.14 | 0.85 | 0.60 | 0.27 |
| V5 and V6 below threshold of .6 |  |  |  |  | 0.33 | 0.17 |  |  |  |
| V5 and V8 below threshold of .6 |  |  |  |  | 0.24 | 1.00 | 1.00 | 0.59 |  |
| V5 and V9 below threshold of .6 |  |  |  |  | 0.43 | 1.00 | 1.00 | 1.00 | 0.20 |
| V5 below threshold of .6 | 0.16 | 0.18 | 0.35 | 0.97 | 0.25 | 0.40 | 0.40 | 0.25 |  |
| V6 and V8 below threshold of .6 |  |  |  |  |  | 0.47 | 1.00 | 0.18 |  |
| V6 and V9 below threshold of .6 |  |  |  |  |  | 0.52 | 1.00 | 1.00 | 0.21 |
| V6 below threshold of .6 | 0.14 | 0.30 | 0.67 | 0.86 | 0.98 | 0.18 | 0.65 | 0.32 | 0.83 |
| V7 and V8 below threshold of .6 |  |  |  |  |  |  | 0.45 | 0.35 |  |
| V7 and V9 below threshold of .6 |  |  |  |  |  |  | 0.57 | 1.00 | 0.27 |
| V7 below threshold of .6 |  |  | 0.47 | 0.76 | 0.97 | 0.97 | 0.29 | 0.23 | 0.12 |
| V8 below threshold of .6 | 0.53 | 0.25 | 0.31 | 0.77 | 0.94 | 0.97 | 0.99 | 0.32 | 0.33 |
| V9 below threshold of .6 | 0.43 | 0.22 | 0.32 | 0.76 | 0.87 | 0.96 | 0.99 | 0.99 | 0.29 |

**Figure S1.**
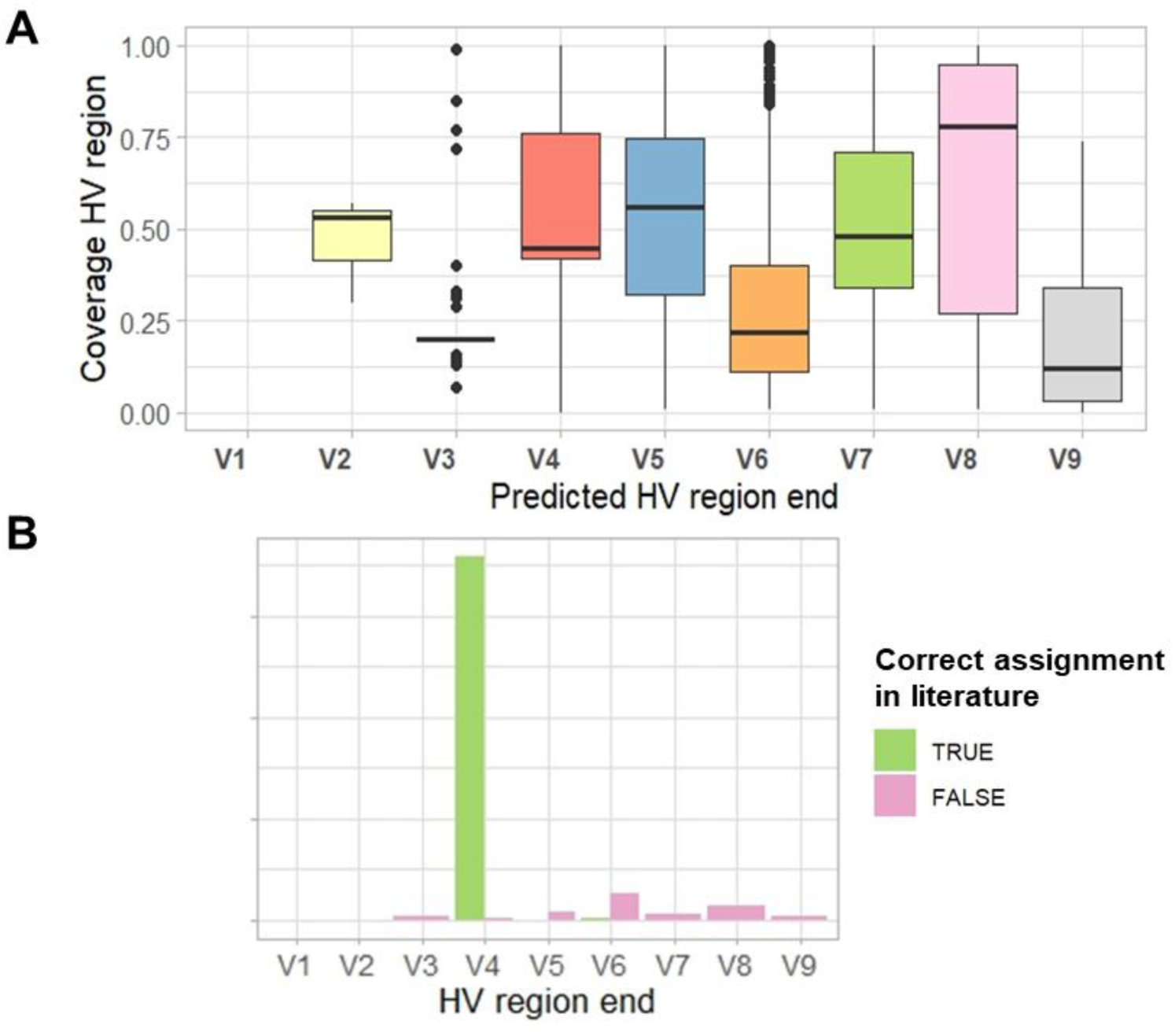
A) Variation in gene coverage across sequences, B) Number of samples in which primer assignment matched the metadata.

## Notes

https://github.com/fbcorrea/hvrlocator

### Competing Interest Statement

The authors have declared no competing interest.

